# Genome-wide and single-base resolution DNA methylomes of the Sea Lamprey (*Petromyzon marinus*) Reveal Gradual Transition of the Genomic Methylation Pattern in Early Vertebrates

**DOI:** 10.1101/033233

**Authors:** Zhao Zhang, Gangbiao Liu, Yangyun Zhou, James P. B. Lloyd, David W. McCauley, Weiming Li, Xun Gu, Zhixi Su

## Abstract

In eukaryotes, cytosine methylation is a primary heritable epigenetic modification of the genome that regulates many cellular processes. While the whole-genome methylation pattern has been generally conserved in different eukaryotic groups, invertebrates and vertebrates exhibit two distinct patterns. Whereas almost all CpG sites are methylated in most vertebrates, with the exception of short unmethylated regions call CpG islands, the most frequent pattern in invertebrate animals is ‘mosaic methylation’, comprising domains of heavily methylated DNA interspersed with domains that are methylation free. The mechanism by which the genome methylation pattern transited from a mosaic to a global pattern and the role of the one or two-round whole-genome duplication in this transition remain largely elusive, partly owing to the lack of methylome data from early vertebrates. In this study, we used the whole-genome bisulfite-sequencing technology to investigate the genome-wide methylation in three tissues (heart, muscle, and sperm) from the sea lamprey, an extant Agarthan vertebrate. Analyses of methylation level and the extent of CpG dinucleotide depletion of gene-encoding, intergenic and promoter regions revealed a gradual increase in the methylation level from invertebrates to vertebrates, with the sea lamprey exhibiting an intermediate position. In addition, the methylation level of the majority of CpGs was intermediate in each sea lamprey tissue, indicating a high level of heterogeneity of methylation status between individual cells. In this regard, we defined the genomic methylation pattern of sea lamprey as “global genomic DNA intermediate methylation”. The methylation features in different genomic regions, such as the transcription start site (TSS) region of the gene body, exon-intron boundaries, transposons, as well as genes grouping with different expression levels, supported the gradual methylation transition hypothesis. We further discussed that the copy number difference in DNA methylation transferases and the loss of the PWWP domain and/or DNTase domain in DNMT3 sub-family enzymes may have contributed to the methylation pattern transition in early vertebrates. These findings demonstrate an intermediate genomic methylation pattern between invertebrates and jawed vertebrates, providing evidence that supports the hypothesis that methylation patterns underwent a gradual transition from invertebrates (mosaic) to vertebrates (global).

## Introduction

Genome-wide cytosine methylation in animals plays important epigenetic roles in modulating gene regulatory networks at different developmental stages and in different tissues (Smith and Meissner 2013). Genome-wide analyses in a number of animal organisms have revealed several interesting features. Global methylation, in which almost all the cytosines in CpG-oligodeoxynucleotides are methylated (except for CpG islands, small unmethylated regions), is the major methylation pattern in jawed vertebrates (Suzuki and Bird 2008; Tweedie et al. 1997). Most of the CpG islands lie in regulatory regions such as promoters, and approximately three-fourths of them are hypomethylated, as required for transcription (Elango and Yi 2008). Moreover, most gene bodies and transposable elements (TEs) in these vertebrates (bony fishes, frogs, birds and mammals) maintain a much higher degree of methylation than flanking regions (Zemach et al. 2010; Feng et al. 2010). In contrast, in invertebrates, including insects (e.g., honey bee, silkworm), echinoderms (e.g., *Echinus esculentus, Strongylocentrotus purpuratus)* and even tunicates (e.g., sea squirt), the genome-wide methylation landscape has revealed a mosaic appearance as the major pattern, in which only specific genomic elements are methylated (Suzuki and Bird 2008; Sarda et al. 2012). More specifically, actively transcribed sequences are hypermethylated. Unlike in vertebrates, TEs in invertebrates are not significantly enriched in methylation compared to adjacent regions (Zemach et al. 2010; Feng et al. 2010). Although the genome-wide distinction of DNA methylation patterns between vertebrates and invertebrates has been well characterized, how and when the transition process has occurred remains largely elusive. Based on the distinct modes between invertebrates and vertebrates, many studies have proposed that such transition might occur rapidly at the time when vertebrates originated (Tweedie et al. 1997; Hendrich and Tweedie 2003; Suzuki and Bird 2008). Alternatively, some studies suggested that the evolutionary transition of the methylation pattern might have involved multiple steps that shaped the different methylation features (Elango and Yi 2008; Okamura et al. 2010). Thus, the transition may have been a complicated process with a long evolutionary span, rather than an immediate, episodic event.

In vertebrates, methylated CpGs that are maintained by conserved DNA methyltransferases are the predominant epigenetic modifications (global methylation), whereas methylated cytosines in CpHpH or CpHpG (H stands for nucleotides A, T or C) constitute only 0.1%-0.2% of the total modifications (Simmen et al. 1999; Ponger and Li 2005; Zemach et al. 2010; Feng et al. 2010). Two key enzymes are DNA methyltransferases and their cofactors (DNMT1, DNMT2, DNMT3, UHRF1 and UHRF2 in mammals) and demethylases (TET1, TET2, TET3 and LSD1 in mammals) (Baubec et al. 2015; Goll and Bestor 2005; Zhang et al. 2011; Albalat et al. 2012; Tahiliani et al. 2009; Shi et al. 2004). Phylogenetic analyses of the DNMT and TET gene families showed that these two gene families expanded in the early stages of vertebrate or chordate evolution (Zemach et al. 2010; Ponger and Li 2005; Albalat et al. 2012; Albalat 2008). It is generally accepted that many genes present in vertebrate genomes owe their origin to one or two rounds whole genome duplications (WGD) that occurred during chordate evolution after the split of the urochordate and cephalochordate lineages but before the radiation of extant gnathostomes (jawed vertebrates) (Wang and Gu, 2002; Smith and Keinath 2015). Hence, an interesting question is how the WGD may have affected the evolutionary transition of the methylation landscape from invertebrates to vertebrates.

The sea lamprey, a modern representative of jawless vertebrates, has been considered an important model species for understanding the origin and diversity of vertebrates. Because of the phylogenetic position of the sea lamprey, a primitive vertebrate positioned between the sea squirt (tunicates) and zebrafish (bony fishes), analyses of its genomic methylation might provide detailed information useful for understanding the origin of vertebrate-specific methylation patterns. In the current study, we used whole-genome bisulfite-sequencing (WGBS) technology to experimentally determine the whole-genome landscape of methylation in three sea lamprey tissues (heart, muscle, and sperm). Together with previous WGBS data, our findings address the following issues: (i) whether the methylation levels of the gene bodies, intergenic regions and transposons differ significantly; (ii) whether the methylation pattern in the sea lamprey is simillar to the canonical pattern in jawed vertebrates; (iii) whether there is a negative correlation between promoter methylation and the methylation of other DNA regions in jawless vertebrates; and (iv) whether the evolution of DNA methyltransferases during the early history of vertebrates could possibly affect the evolution of vertebrate-specific methylation patterns. Here we describe a general picture of the dynamic methylome evolution, which characterizes the transition process from invertebrate-specific to vertebrate-specific methylation patterns.

## Results

### Landscape of genome-wide DNA methylation pattern in sea lamprey

We obtained whole-methylome bisulfite sequencing data at single-base resolution from three tissues (heart, muscle and sperm) of a male sea lamprey. We also extracted corresponding total RNA from the heart sample and performed RNA sequencing. The annotated sea lamprey genome consisted of approximately 0.6 Gbp, including 27.5 million CpG sites (CGs) (single strand). Genetic polymorphisms (single-nucleotide polymorphisms, SNPs) could potentially disrupt the calling of the methylation status of cytosines. Therefore, we performed whole-genome resequencing of genomic DNA extracted from the heart tissue (5X coverage) and identified approximately 0.18 million SNPs between our sample and the reference genome (version *Pmarinus_7.0).* Indeed, approximately 9,000 CGs were disrupted by SNPs in our samples (<0.01%). Thus, these sites were excluded from further analyses. To estimate the efficiency and specificity of our bisulfite conversion, we spiked unmethylated lambda DNA and methylated human DNA standard into each library as negative and positive control, respectively (see Materials and Methods). On average, fewer than 0.2% of the unmethylated cytosines in lambda DNA were estimated to fail to undergo the C–T conversion during bisulfite treatment for each library, and fewer than 5% of the methylated cytosines in CGs of positive control were estimated to be over converted.

We sequenced approximately 140.12, 60.18 and 65.07 Gbp of WGBS data for the heart, muscle and sperm samples, respectively. We aligned bisulfite-concerted reads against the reference genome (version *Pmarinus_7.0)* using the Bismark program (Krueger and Andrews 2011). After data preprocessing, approximately 39% of the resulting reads were uniquely mapped to the sea lamprey reference genome. We achieved 20X, 9X and 10X coverage per strand for the heart, muscle and sperm, respectively. Approximately 81.3%, 38.6% and 19.3% of the genomic cytosine were covered by at least two unique reads in the heart, muscle and sperm samples (supplementary table 1), respectively. In total, over fifty million cytosines in CpGs were covered, and more than 21 million cytosines were shared among the three tissues (supplementary figure 1A). Nearly 10 million of the shared CpGs were covered by at least 5 reads (supplementary figure 1B). The relatively low mapping rate of our samples is probably attributable to the incompleteness of the sea lamprey draft genome, which are mainly due to the effects of programmed genome rearrangement, and highly repetitive, high GC content and highly heterozygous of the sea lamprey genome (Smith et al. 2013).

We calculated the methylation level of each cytosine in the CpG, CpHpG and CpHpH sites and found a similar distribution among the three tissues (figure 1A). On average, approximately 51% (50.9%, 50.4% and 50.7% for heart, muscle and sperm, respectively) of the sequenced CGs were methylated (coverage >5 reads, FDR < 0.01) (see Materials and Methods). Among all the methylated cytosines (mCs), more than 99% were in CpGs (supplementary table 1), which is consistent with previous results in other animal methylomes (Zemach et al. 2010; Feng et al. 2010). Most of the methylated CpGs (mCGs) exhibited intermediate methylation levels (between 10% and 90%, figure 1A), which is inconsistent with the high methylation level observed in other vertebrates (Zemach et al. 2010; Feng et al. 2010). This observation suggested that the methylation status of most CGs in sea lamprey is highly variable between individual cells. We did not observe significant methylation at non-CpG sites in any of the samples (figure 1B, supplementary table 2). Therefore, all subsequent analyses were focused mainly on the CpG sites.

**Figure 1.**
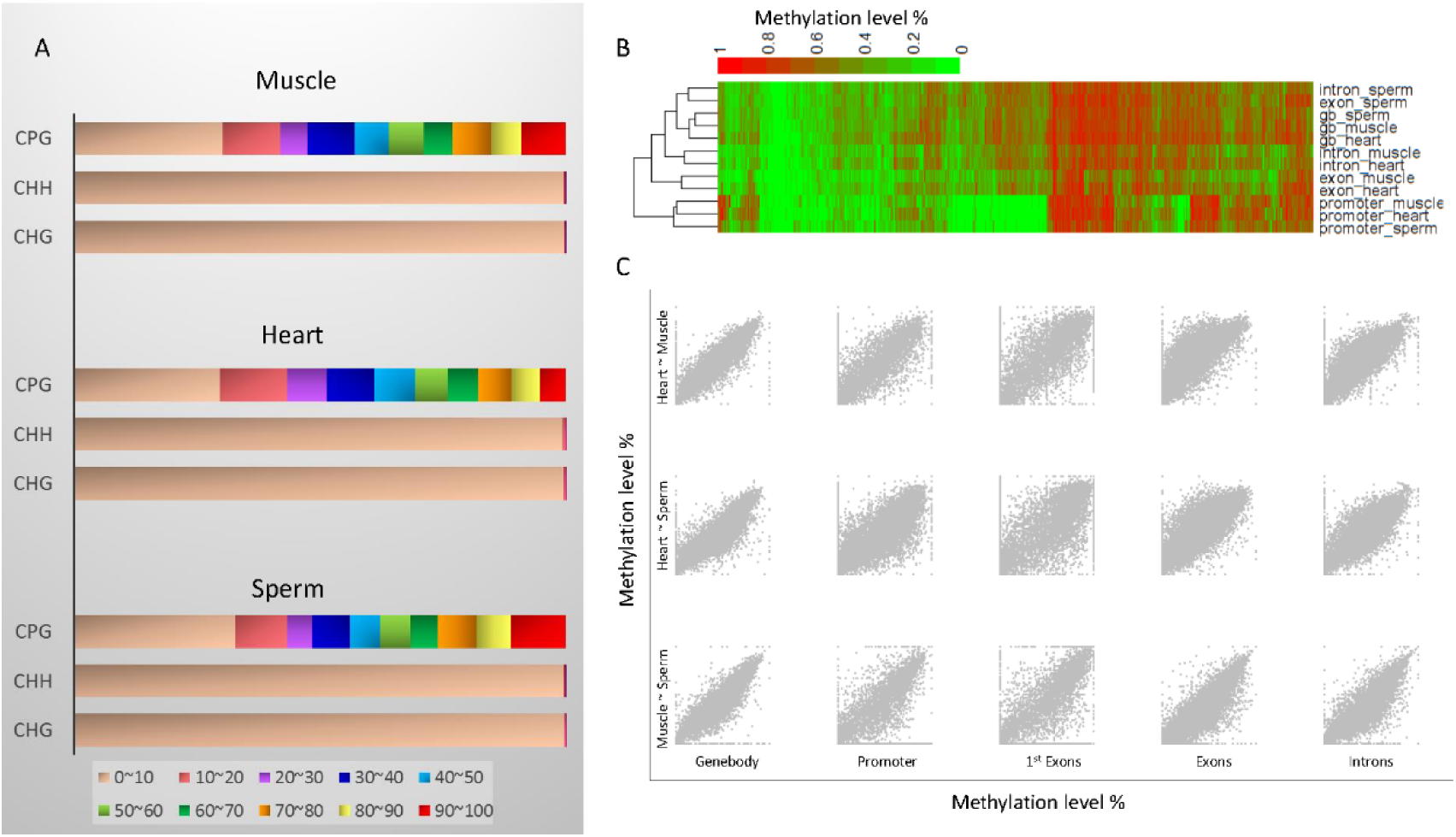
Comparison of the genome methylome of sea lamprey muscle, heart and sperm. A) The methylation level and distribution of methylated C in CpG, CHH, and CHG regions for the three tissues. B) Heat map showing the clustering of CpG methylation levels in the gene bodies, exons, introns and promoters of the three tissues. C) Pairwise comparison of the gene bodies, promoters, exons, introns, and the 1st exons among the three tissues.

Next, we examined the methylation features in various functional elements such as promoters, gene bodies, exons, and introns. The results showed that the unmethylated CGs were highly enriched in promoters (figure 1B). Unsupervised clustering revealed that the methylation similarity among the three tissues was higher in the promoter and gene body than in introns and exons (figure 1B). The levels of CpG methylation in specified regions of the three tissues were consistent (figure 1C). In general, the wholegenome methylation pattern in the three tissues was largely similar, so the heart tissue, which had the highest WGBS sequencing coverage, could be considered representative to explore DNA methylation patterns in sea lamprey in more depth.

### CpG O/E ratio and methylation level distributions in promoters and gene bodies are gradually transited from invertebrates to vertebrates

Most DNA methylation in vertebrate genomes occurs at CpGs. Methylated cytosines are hypermutable due to their vulnerability to spontaneous deamination, which results in a gradual depletion of CpG dinucleotides from methylated regions over time. Thus, the CpG O/E ratio (CpG_O/E_) has been widely used as a robust indicator of the level of historical germline DNA methylation in a specified chromosomal region (such as Elango and Yi 2008; Weber et al. 2007; Suzuki et al. 2007). Briefly, a low CpG_O/E_ indicates that the specific region may have undergone germline hypermethylation, whereas a high ratio reflects germline hypomethylation (Suzuki and Bird 2008). For example, according to the CpG_O/E_, human promoters can be divided into high CpG_O/E_ promoters (CpG_O/E_ above 0.75) and low CpG_O/E_ promoters (CpG_O/E_ below 0.5), which tend to be hypomethylated or hypermethylated, respectively (Weber et al. 2007). Previous studies have reported a bimodal CpG_O/E_ distribution for gene bodies and intergenic regions in invertebrates based on the mosaic genomic methylation pattern (Elango and Yi 2008; Okamura et al. 2010). Thus, some genomic regions are hypermethylated and some are hypomethylated (Elango and Yi 2008; Weber et al. 2007; Suzuki et al. 2007). The CpG_O/E_ of promoters in invertebrates are unimodal, indicating a uniform low methylation pattern. In contrast, vertebrate gene bodies and intergenic regions exhibit a bimodal distribution, and vertebrate promoters show a bimodal CpG_O/E_ distribution, suggesting a high level of methylation throughout the genome, except a small portion of the genome with low methylation in promoter regions. The bimodal CpG_O/E_ distribution of vertebrate promoters was speculated to associate with development and gene expression regulation (Tweedie et al. 1997; Elango and Yi 2008; Sarda et al. 2012; Weber et al. 2007; Suzuki et al. 2007; Okamura et al. 2011).

However, whether the transition from the invertebrate mosaic DNA methylation pattern to the vertebrate whole-genome methylation pattern occurred gradually or episodically remains unclear. To address this issue, we estimated the CpGo/e in gene bodies and promoters in seven animals, including two invertebrates: honey bee *(Apis mellifera)* and sea squirt *(Ciona intestinalis);* one jawless vertebrate: sea lamprey *(Petromyzon marinus);* one cartilaginous fish: elephant shark *(Callorhinchus milii*); one bony fish: zebrafish *(Danio rerio);* one amphibian: western clawed frog *(Xenopus tropicalis);* and one mammal: human *(Homo sapiens)* (figure 2A). Interestingly, the CpG O/E ratios of the sea lamprey promoter and gene body were both unimodal, and the values were between those of invertebrates and vertebrates, suggesting that the whole genome methylation pattern in sea lamprey is not ‘mosaic’ and the whole methylation level is between known invertebrates and vertebrates. The median CpG_O/E_ of the sea lamprey gene body/promoter was 0.69/0.89, which were lower than those determined for honey bee (1.10/1.39) or sea squirt (0.80/0.91) but greater than those for zebrafish (0.50/0.61) or human (0.41/0.51). Moreover, we calculated the methylation level of the gene bodies and intergenic regions in five animals (honey bee, sea squirt, sea lamprey, zebrafish and human) and discovered that the methylation level gradually increased from invertebrates to vertebrates (supplementary table 2). Sea lamprey genomic DNA undergoes programmed genome rearrangement during embryonic development, which leads to systematic gene loss and may have biased the results of our methylation analyses (Smith et al. 2012). To eliminate this potential influence, we selected genes with germline-specific expression in isolation to detect the CpG_O/E_ for gene bodies, promoters, exons and introns using the aforementioned method and obtained results consistent with those from the somatic tissues (supplementary figure 2).

**Figure 2.**
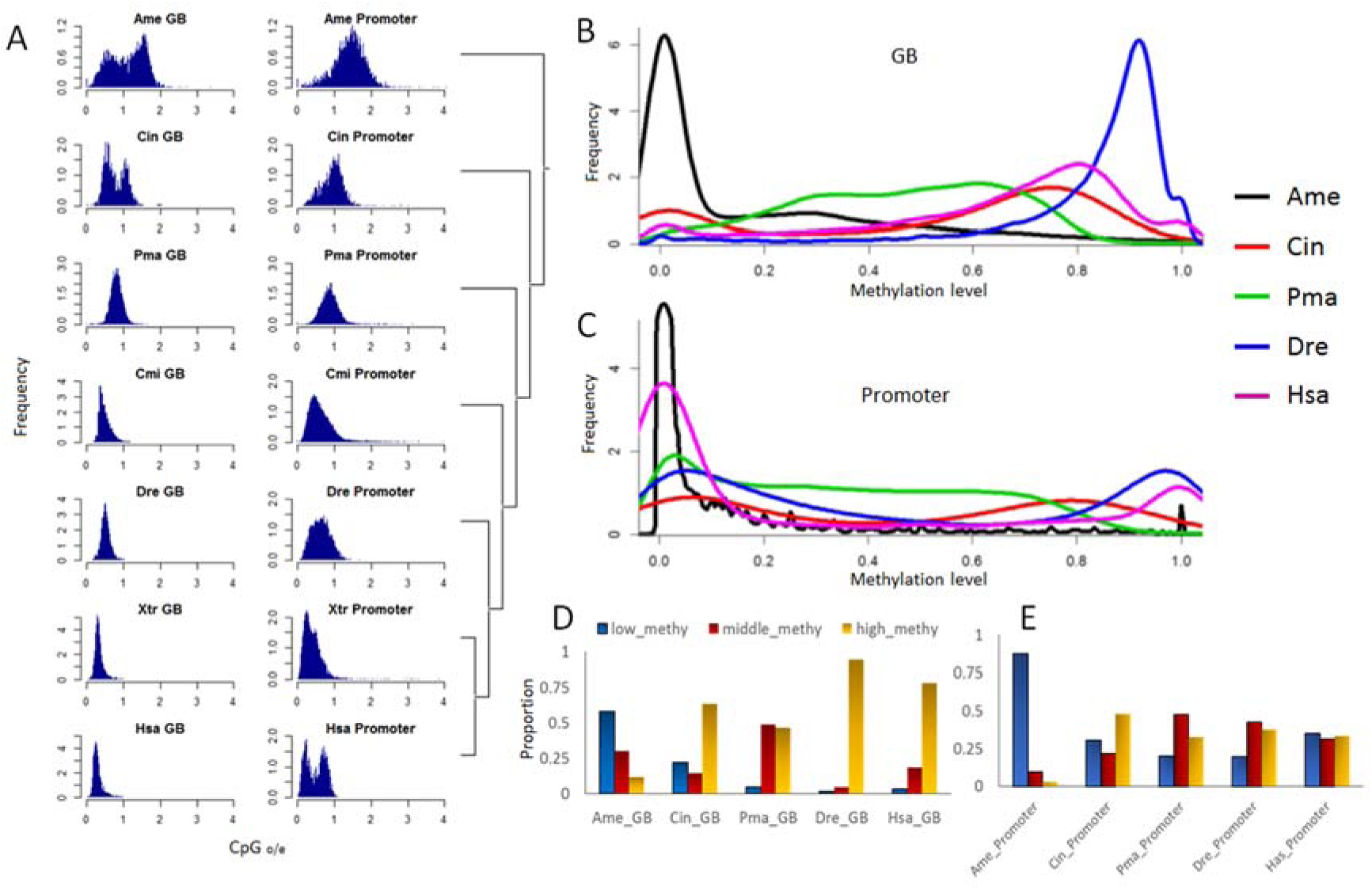
Landscape of animal methylation. A) The distribution of the CpG_O/E_ of the gene bodies and promoters. B) and C) Distribution of the methylation level in gene bodies and promoters, respectively. D) and E) Proportion of gene bodies and promoters classified in the high methylation, intermediate methylation, and low methylation groups. Ame: honey bee; Cin: sea squirt; Pma; sea lamprey; Cmi: elephant shark; Dre: zebrafish; Xtr: claw frog; Hsa: human.

To confirm the results obtained for the CpG_O/E_, we detected the methylation level for the gene bodies of protein-coding genes and their promoters in vertebrates and invertebrates (figure. 2B and figure. 2C). Consistent with the bimodal CpG_O/E_ pattern, the methylation of gene bodies diverged into hypermethylation and hypomethylation in invertebrates (sea squirt and honey bee), whereas they were almost uniformly hypermethylated in vertebrates. Interestingly, an intermediate distribution of the methylation level was identified in sea lamprey genes (from 0.1 to 0.9, without an obvious peak). As previously reported, promoters are mainly hypomethylated in the honey bee but are divergent in sea squirt and vertebrates. We also defined three methylation level categories, low (<0.1), intermediate (>0.1 and <0.5), and high (>0.5), to quantify the methylation transition for the gene bodies and promoters in the five species (figure 2D and figure 2E). We found that most gene bodies of sea lamprey were intermediately methylated, in contrast to invertebrates (dominated by low methylation) or other vertebrates (high methylation). These findings indicated that the methylation pattern in lamprey differed from known patterns (mosaic and global methylation in invertebrates and vertebrates, respectively).

From invertebrates to vertebrates, we found that the proportion of gene bodies with low methylation decreased and that those with intermediate to high methylation increased (figure 2D). The majority of the promoters in honey bee had a low level of methylation. Differences in the methylation of promoters in vertebrates were not significant (figure 2E). In summary, these findings revealed that the CpG_O/E_ distribution gradually changed and that the methylation level in gene bodies and promoters steadily increased during early vertebrate evolution.

### Global intermediate methylation pattern in sea lamprey genomic DNA

The whole-genome methylation pattern is mosaic in invertebrates and global in higher vertebrates (teleosts and tetrapods). How these patterns evolved is still unclear. In the present study, we obtained the sea lamprey methylome to infer the evolution of the genomic methylation pattern in early vertebrates. We analyzed the methylation level of whole-genome CpGs and found that sites with low methylation (<0.1) were predominant in sea squirt (75%) whereas those with high methylation (>0.5) were prevalent in zebrafish and human (~82% and ~76%). However, most CpGs were intermediately methylated in lamprey (<0.5 and >0.1, 46%) (supplementary figure 3B). We randomly selected Chr1 from human, zebrafish, sea squirt and scaffold GL4763991 from lamprey as an example to display the methylation level of CpGs (figure 3A). As shown in the figure, a global high methylation pattern was detected in zebrafish and human, and a mosaic pattern was identified in the sea squirt. Interestingly, we found that the sea lamprey genome was also characterized by global methylation, with the majority of the CpG sites having intermediate methylation level, which means a medium percentage of cells bearing a methylated cytosine at a given locus. To confirm this observation, we examined other chromosomes (scaffolds) using the same method and found a consistent pattern in the majority of cases (supplementary figure. 3A). Additionally, we selected a conservation of syntenic chromosome regions with one-to-one orthologous genes (NDUFB5, MPRL47) to compare the methylation pattern in the sea squirt, sea lamprey, zebrafish and human genomes. The results were similar to those determined for randomly selected regions (figure 3B).

**Figure 3.**
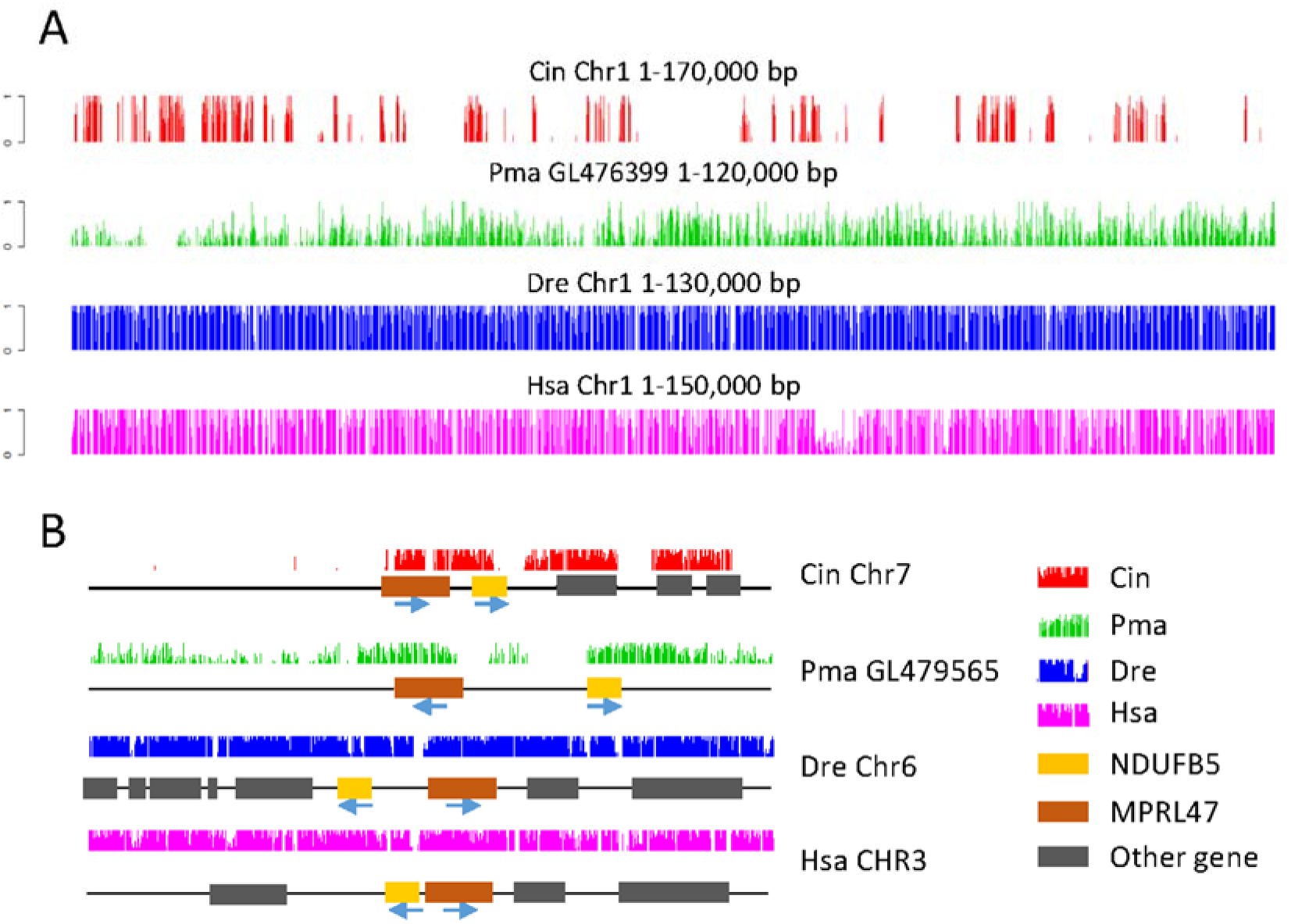
Genome browser representations of DNA methylation patterns. A) DNA methylation pattern for randomly selected genomic regions in the sea squirt, sea lamprey, zebrafish and human. B) DNA methylation pattern for a homologous chromosome segment in the four animals. Yellow and grown squares stand for the two homologous genes (NDUFB5 and MPRL47) and gray squares represent other flanking genes. Abbreviations are the same as in figure 2.

To further validate the global but not mosaic genome methylation pattern in sea lamprey, we then examined the genome-wide distribution of average methylation level for sea squirt, sea lamprey and zebrafish genome, with a sliding window of 10kb or 1kb (supplementary figure 4). We found a bimodal distribution in sea squirt, representing comparable amounts of low- and high-methylation-level regions, a mosaic genome methylation pattern (supplementary figure 4). While in sea lamprey and zebrafish, only one intermediate- or high-methylation-level peak was conspicuous, reflecting a global methylation pattern (supplementary figure 4B-C). Moreover, there are many methylation free short regions (1kb) in sea squirt and sea lamprey genomes, but not in zebrafish genome (supplementary figure 4D-F). These results indicate that, in sea squirt genome, most consecutive CpGs were unmethylated, while in lamprey and zebrafish, unmethylated CpGs were rare. The peak of average methylation level distribution for each region in sea lamprey (0.34) was lower than in zebrafish (0.93), which suggested the methylation level of sea lamprey was intermediate between sea squirt and zebrafish. Hence, combined the results of methylation level and the extent of CpG dinucleotide depletion analyses, we defined the lamprey genome methylation pattern as “global intermediate methylation”, as opposed to the “global high methylation” pattern observed in vertebrates and the “mosaic methylation” pattern identified in invertebrates. These results support our hypothesis that the whole-genome methylation pattern evolved gradually in early vertebrates.

### Methylation features of the sea lamprey genome: transition state from invertebrates to vertebrates

To further characterize the genome methylation pattern transition in early vertebrates, we analyzed several methylation features in honey bee, sea squirt, sea lamprey, zebrafish and human. First, we calculated the relative difference in CpG methylation for all pairwise species, which reflects the methylation level divergence in specific regions (see Materials and Methods). We found that the difference between lamprey and zebrafish was similar to that between lamprey and sea squirt in both the gene body and the promoter (0.156–0.18 for the gene body, 0.036–0.072 for the promoter, supplementary table 3). Second, we profiled the DNA methylation levels across the gene bodies. Our data showed that CpG methylation exhibited a characteristic peak in the bodies of protein-coding genes in honey bee, sea squirt and sea lamprey, whereas in zebrafish and human, the methylation level in the gene bodies was similar to that in the genome background (figure 4). The methylation level in the promoter region (the region around the TSS) in sea squirt and lamprey was slightly decreased compared with the intergenic region, but the methylation level in the promoter region in zebrafish and human was greatly decreased compared with the surrounding region (figure 4). Sea lamprey and sea squirt showed similar decreases in methylation level in the transcript end site (TES) region, whereas zebrafish and human had no obvious change. In addition to CpG methylation, we also analyzed the CG dinucleotide content of the gene bodies and found that the features of lamprey CG content were also more similar to sea squirt than to other vertebrates (supplementary figure 5). Additionally, using one-to-one orthologous genes with conserved synteny in the four chordates (sea squirt, sea lamprey, zebrafish, human), we analyzed the methylation level of some syntenic regions and found that the gene body/promoter methylation levels in different species were similar at the level of individual genes (supplementary figure 6).

**Figure 4.**
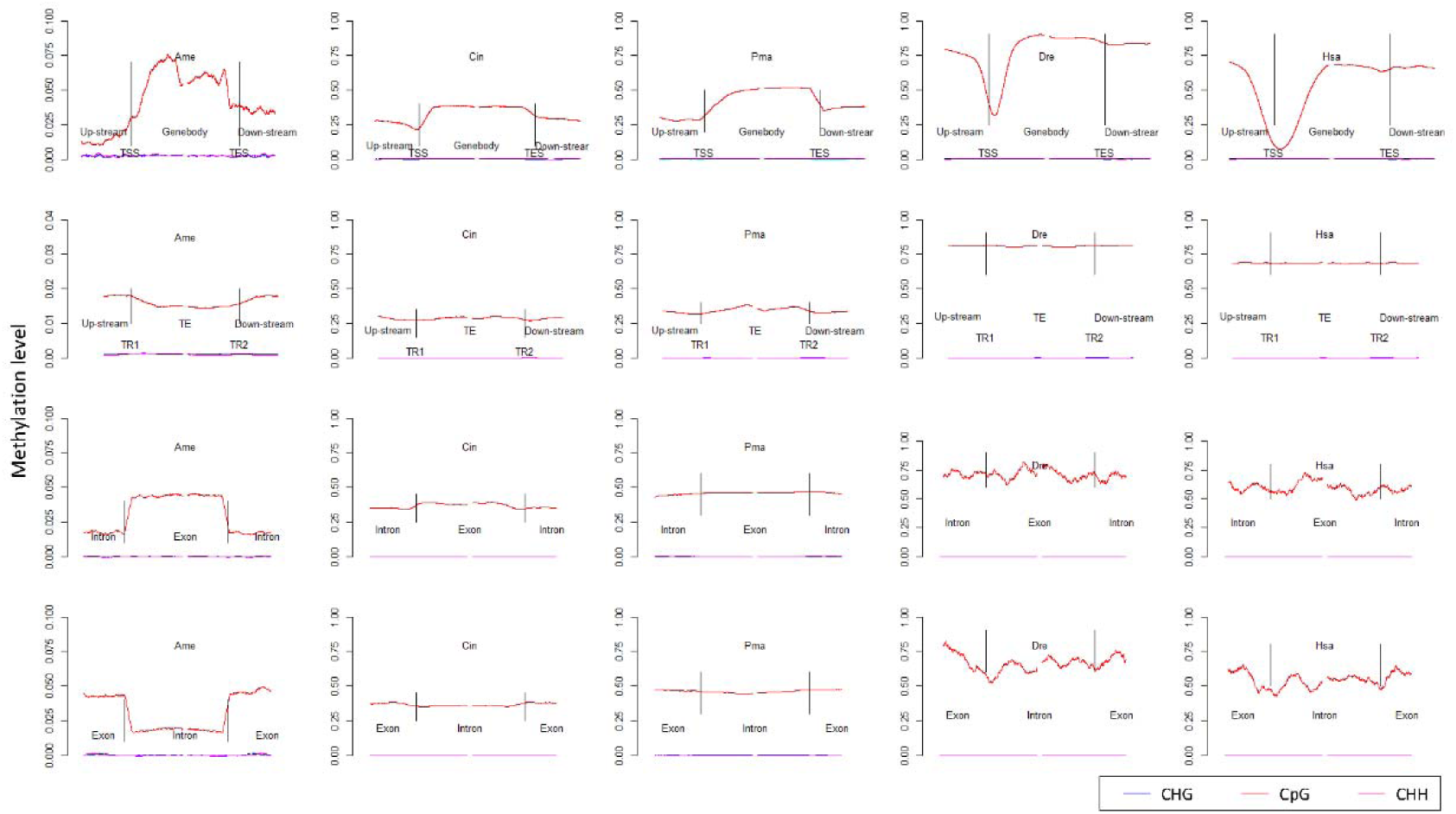
DNA methylation in different genomic regions in the honey bee, sea squirt, sea lamprey, zebrafish and human. Only data obtained for up to halfway to the next gene body, transposon, exon or intron were used in this analysis. The gene bodies and transposons were aligned at the 5 ‘ end (left vertical line) or the 3’ end (right vertical line). The average methylation levels for each 300-bp, 200-bp, 20-bp and 20-bp interval were plotted for the gene bodies, transposons, exons and introns, respectively. The upstream region, body region, and downstream region of the gene body were 4 kb, and those for transposons were 3 kb. The average methylation levels for each 20-bp interval were plotted in the exons and introns. The upstream region, exons/introns and downstream region for the exons/introns were 200 bp. TE: transposon element; TR: transposon repeat; TSS: transcript start site; TES: transcript end site. Other abbreviations are the same as in figure 2.

We also extracted multi-exon genes to analyze the methylation level in exon-intron-exon structures (figure 4). Honey bee and sea squirt exons had obviously higher CpG methylation levels than introns. In contrast, in sea lamprey, zebrafish and human, exons and introns shared similar methylation levels. Next, we compared the methylation levels of transposons in the five animals. In the two transposon repeat regions, we divided the upstream, transposon, and downstream regions and calculated the methylation level. In honey bee transposons, CpGs were clearly hypomethylated (figure 4). In sea squirt and sea lamprey, CpG methylation in transposons was higher than that in the flanking regions; however, in zebrafish and human, the methylation level in transposons was similar to that in surrounding regions. In summary, these results suggest that the vertebrate-specific methylation features may not have matured yet in the lamprey, or that the sea lamprey lineage lost some vertebrate-specific methylation features.

### Methylation, CG content and expression of transcribed genes

DNA methylation plays an essential role in the regulation of gene expression. To understand the influence of methylation on the regulation of gene expression in sea lamprey, we analyzed the methylation level for groups with different expression levels in the sea squirt, sea lamprey and zebrafish. The sea lamprey DNA methylation and gene expression data were obtained from the heart tissue, and those for sea squirt and zebrafish were derived from previous RNA-seq studies (see Materials and Methods). We found that ~35% of the genes were not expressed in lamprey heart (FPKM = 0). By ranking the gene expression level, we divided the genes into five groups from high to low in each species as follows: 1st (the 10% highest-expressed genes), 3rd (10%-30%), 5th (30%-50%), 6th (50%-65%), and 10th (65%-100%). The DNA methylation pattern and gene expression in the three species exhibited two consistent features: 1) the promoter region of the highly expressed genes had a low level of methylation; in contrast, the promoter region of genes with a low expression level was highly methylated; 2) the bodies of low-expression genes had a low level of methylation (figure 5A-C). This pattern is similar to that in Arabdiopsis (Coleman-Derr et al. 2012), suggesting a similar role of DNA methylation on gene expression regulation between plants and animals. Additionally, we analyzed the GC content of each gene category and discovered the following: 1) genes with low expression displayed a low level of GC content in the gene body (figure 5D-F); 2) the TSS and transcription end site (TES) regions were CpG-poor or CpG-dense for genes with low or high expression, respectively.

**Figure 5.**
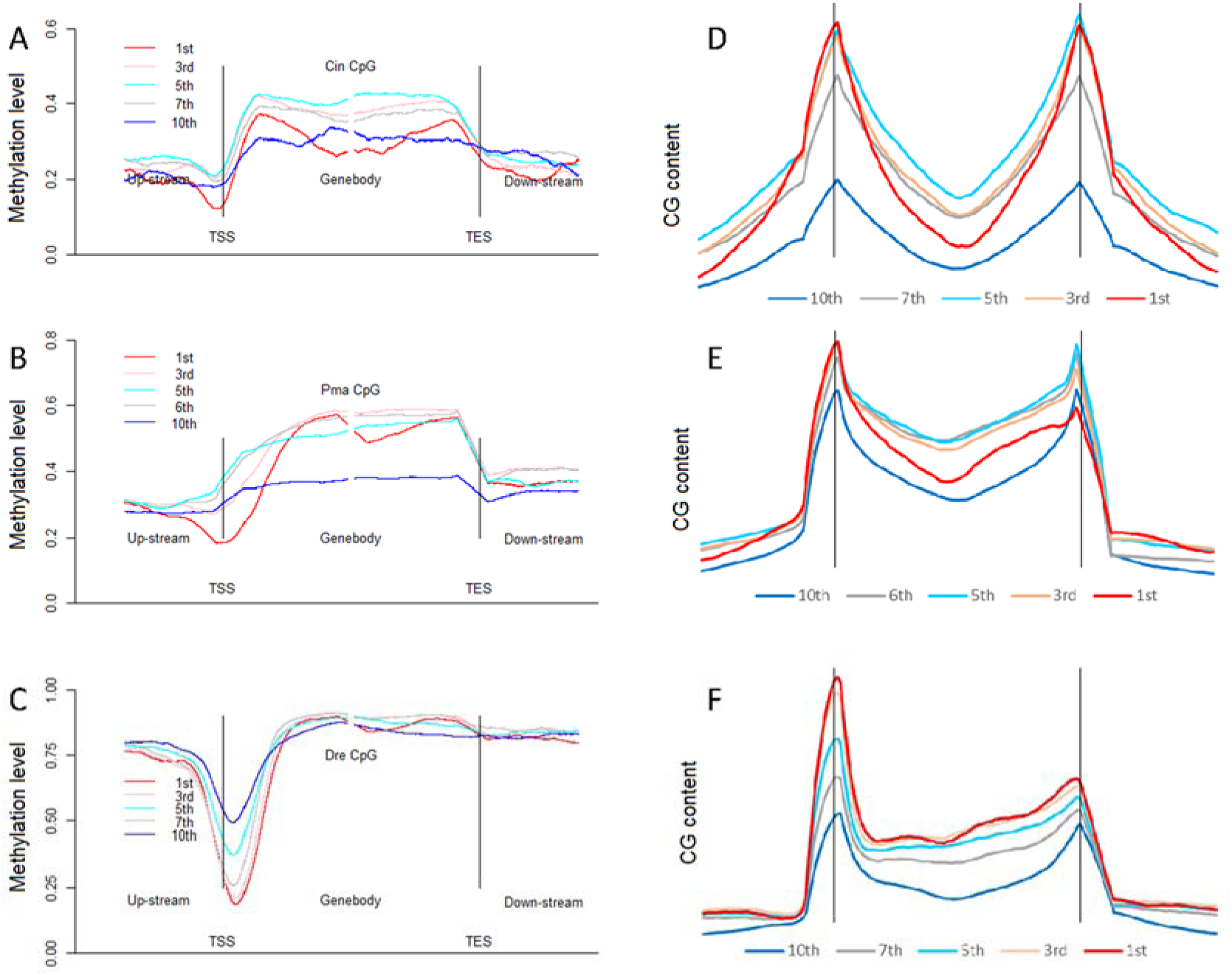
Distribution of CpG methylation and GC content for protein-coding gene groups with different expression levels. A-C) CpG methylation pattern in different expression groups. D-F) GC content in different expression groups. Abbreviations are the same as in figure 2 and figure 4.

### Phylogenetic analysis of the DNA methylation machinery

The establishment of DNA methylation patterns is mediated by DNA methyltransferases and the enzymes involved in demethylation, especially DNMT3A and DNMT3B, which bind to the chromosome and mediate de novo DNA methylation (Cheng and Blumenthal 2008). Dnmt3L, a cofactor for de novo methyltransferases, stimulates Dnmt3A/3B activity by forming the DNMT3A-DNMT3L complex (Jia et al. 2007; Gowher et al. 2005). DNMT3A and DNMT3B have a similar domain arrangement and perform similar functions in the genomic methylation process (Baubec et al. 2015; Cheng and Blumenthal 2008). The DNMT3 proteins have two important domains for its function: the PWWP domain, which can non-specifically bind DNA, and the C-terminal catalytic domain (Baubec et al. 2015; Cheng and Blumenthal 2008; Qin and Min 2014) (also known as the DNA MTase or DNA methylase domain), which can catalyze DNA methylation and mediate the formation of the 3A-3A or the 3A-3L complex (Kato et al. 2007; Li et al. 2007; Van Emburgh and Robertson 2011). To explore the potential mechanisms underlying the formation of sea lamprey and vertebrate-specific methylation patterns, we performed phylogenetic analyses of several methylation-related proteins. We found that the copy number of DNMT3 genes changed in different lineages; for example, DNMT3B was not present in sea squirt or lamprey (figure 7A, supplementary figure 7B). Additionally, the functional domain of DNMT3 in lamprey differed from that in other vertebrates: 1) DNMT3Ab in lamprey had no PWWP domain, which was present in all DNMT3s in zebrafish and human; 2) lamprey DNMT3s contained only a single DNA methylase domain, whereas all DNMT3s in zebrafish and human had two copies of the DNA methylase domain (figure 6B). This results led us to further investigate the domain composition of DNMT3s chicken, which has been reported to own significantly lower genome DNA methylation level than zebrafish and human (Mugal et al, 2015). Interestingly, we found chicken DNMTs also contained a single DNA methylase domain. In addition, we investigated several other DNA methyltransferases, such as DNMT1, that ensure the maintenance of methylation through replication, and we discovered a KG-enrichment motif in vertebrate DNMT1 (supplementary figure 7A, C, D). Other DNA methylases, such as DNMT2, UHRF1, and UHRF2, and demethylases (TET1, TET2, and TET3) were identical in both gene copy number and domain constitution among analyzed species (supplementary table 4). Together, these results suggest that DNA methyhransferases, rather than other methylases, may play an important role in CpG methylation pattern transition in early vertebrates.

**Figure 6.**
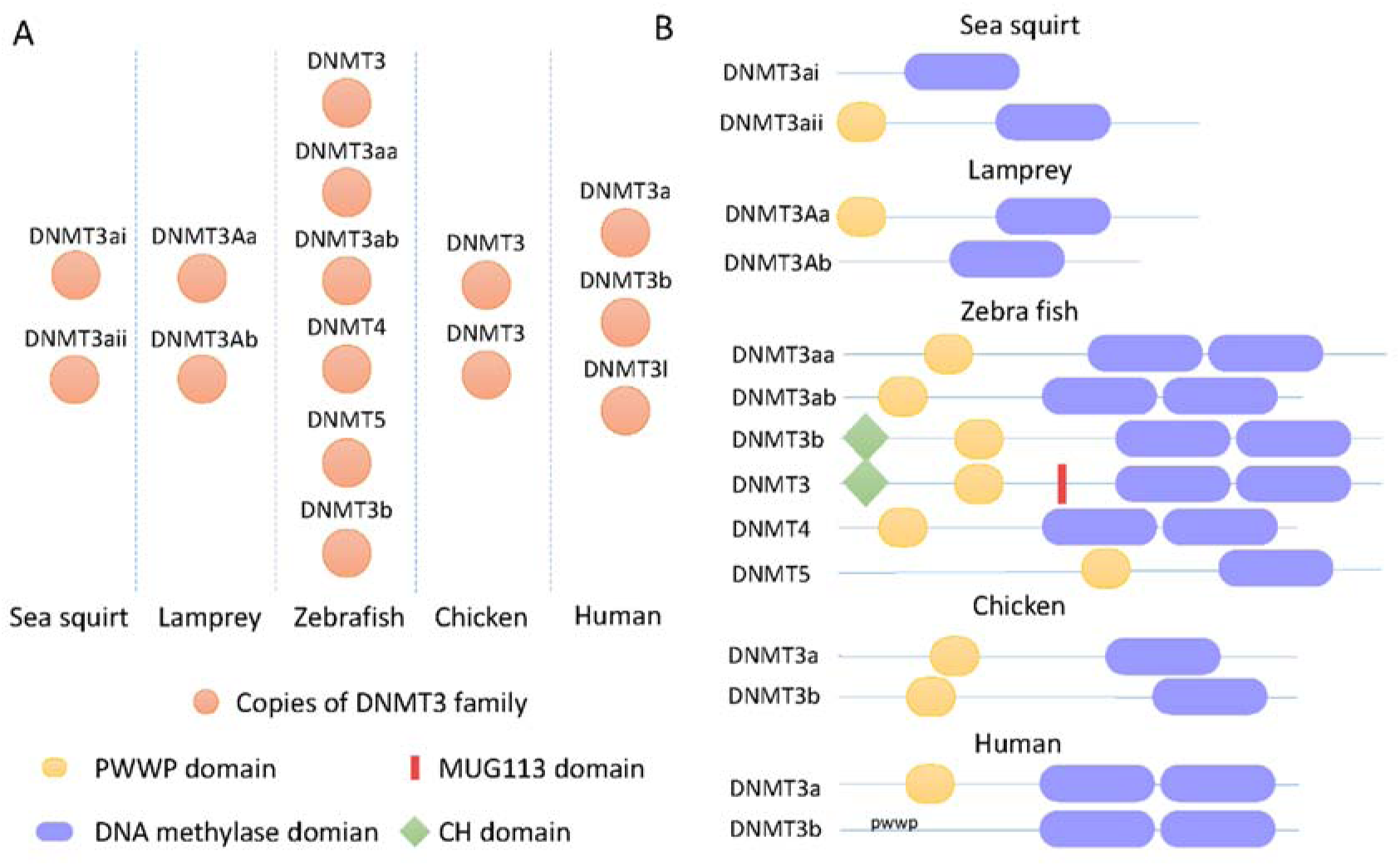
Copy number and domain constitution of the DNMT3 sub-family in sea squirt, sea lamprey, zebrafish, chicken and human. A) Copy number of DNMT3s. B) Domain constitution of DNMT3s.

## Discussion

CpG methylation is an important mechanism in the regulation of gene expression and in many important cellular processes in both vertebrates and invertebrates. Divergent methylation patterns exist between invertebrates (e.g., inserts, tunicates, amphioxus (Albalat 2008; Elango et al. 2009; Wang et al. 2006; Suzuki et al. 2013)) and vertebrates (e.g., bony fishes, tetrapods (Suzuki and Bird 2008; Elango and Yi 2008; Albalat et al. 2012)). The divergence of species from the most recent common ancestor of vertebrates and tunicates occurred ~750 million years ago, and bony fishes emerged more than ~450 million years ago (Blair and Kumar 2003). Although more than 20 eukaryotic methylomes have been examined to date, methylome data are still lacking for species positioned between the lineage of tunicates and bony fish, which span approximately 300 million years. Thus, the transition in the whole-genome methylation pattern during this stage remains unclear. Many studies have compared the CpGo/e and methylation level of gene bodies and promoters to define a “gradual genomic DNA methylation transition” hypothesis from a mosaic methylation pattern in invertebrates to a global methylation pattern in vertebrates (i.e. Weber et al. 2007; Suzuki et al. 2007; Elango and Yi 2008). However, until now, there has been a lack of direct evidence for this hypothesis (Okamura et al. 2010). The jawless fishes represent one transition lineage between tunicates and bony fishes (Lagman et al. 2013; Mehta et al. 2013; Decatur et al. 2013; Donoghue and Purnell 2005). As a jawless fish with an assembled genome, the sea lamprey genome methylation pattern may explain how the pattern transitioned from mosaic to global genomic DNA methylation.

In this study, we conducted both CpG_O/E_ based DNA methylation level estimation and whole genome bisulfite sequencing to test this hypothesis. The CpG_O/E_ has been widely considered as a well established signature of the historical germline DNA methylation level in a specified chromosomal region. We compared the landscape of five species (including two invertebrates, honey bee and sea squirt, and three vertebrates, sea lamprey, zebrafish and human) to estimate the normalized CpG_O/E_ distribution in gene bodies and promoters. Our results revealed that the distribution of the CpG_O/E_ was unimodal in both the lamprey gene body and promoter, suggesting that the genomic DNA methylation pattern in lamprey may differ from other known species. The prediction of the CpG_O/E_ was further validated by our WGBS data. The consistence of the results of CpG_O/E_ analysis and WGBS experiments revealed that CpG_O/E_ could be considered as a robust indicator for evolution of methylation.

The single-base resolution methylomes of three tissues of sea lamprey provided a more precise DNA methylation level estimation than CpG_O/E_ analysis. In our WGBS experiments, we spiked in both negative and positive controls and have shown the highly efficient conversion of unmethylated cytosines (false-positive rate < 0.2%) and the low over-conversion rate (false-negative rate < 5%) for the methylated cytosines. It is notable that CpG sites in the positive control are not completely methylated, as indicated by the manufacturer. So the real false-negative rate should be smaller than our observation. Additionally, the results from three different libraries are consistent. Together these results demonstrated our results are highly reliable. We found that the methylated regions in sea lamprey were consecutive; however, the methylation level was lower than in other vertebrates. To distinguish this feature from known methylation patterns, we defined the lamprey pattern as “global genomic DNA intermediate methylation”. We also analyzed the methylation characteristics in selected species and found 1) the genome methylation pattern divergence extent between sea lamprey and sea squirt or other methylome available vertebrates are almost equivalent; 2) the methylation level in the promoter region in sea lamprey demonstrated a greater similarity to sea squirt than to other vertebrates (for example, in the promoter region, the decreased methylation detected in sea squirt/lamprey was not apparent in other vertebrates), but the inverse situation was observed in the transcript end site (TES) region; 3) the difference in the intragenic methylation level (exon-intron-exon or intron-exon-intron) was reduced in lamprey and vertebrates compared to invertebrates; and 4) transposon methylation in lamprey demonstrated a greater similarity to that in sea squirt than in vertebrates. These observations reveal that the methylation features in lamprey represents an intermediate status between invertebrates and vertebrates.

Divergent DNA methylation patterns might be attributed to the divergence of DNA methyltransferases in different eukaryotes. The DNMT3 subfamily performs *de novo* methylation, which may significantly influence the genomic DNA methylation pattern (Baubec et al. 2015; Jia et al. 2007; Gowher et al. 2005; Qin and Min 2014; Kato et al. 2007; Li et al. 2007; Van Emburgh and Robertson 2011; Yan et al. 2011). We analyzed the copy numbers and constitution of DNMT3 proteins in sea squirt, zebrafish and human. We were unable to identify the DNMT3B gene in the sea squirt or sea lamprey genome, so these two species may have lost the DNMT3B gene independently. Further research is needed to confirm the loss of the DNMT3B gene in the sea squirt and sea lamprey genomes, instead of being undetectable due to incomplete genome sequencing data. Additionally, two important domains involved in *de novo* DNA methylation, the PWWP domain and DNA MTase, were incomplete in sea lamprey (PWWP domain loss or DNA MTase deficiency in one copy of lamprey DNMT3A). This phenomenon may have contributed to the non-canonical global DNA methylation pattern in early vertebrates. Interestingly, we found that the DNMT3 copy number and domain composition in birds and reptiles are similar to sea lamprey. For example, there are two DNMT3 genes in chicken genome, and each DNMT3 gene contains one methylase domain. According to a recent study, whole genome DNA methylation level of chicken sperm is about 41%, which is very similar to sea lamprey (Mugal et al. 2015). It will be of interest to investigate whether the genome methylation pattern is similar between sea lamprey and chicken or other reptiles, such as lizard and turtle. Together, these findings suggest that the evolution of DNMT family in vertebrates possibly contributed to the genomic methylation pattern change in vertebrates. Furthermore, environmental influence during vertebrate evolution may also have contributed to the global DNA methylation change. For example, it has been reported that the whole genome methylation level in reptiles and fishes are negatively correlated with habitat temperature, independently of phylogenetic distance (Varriale, 2014). It will be interesting to test if the environment has influence on evolutionarily genome methylation pattern transition in the early vertebrates.

From an evolutionary perspective, during the transition from invertebrate to vertebrate, both the 1R/2R whole genome duplication and advent of epigenetic regulations, including DNA methylation, may have contributed to the evolution of complexity of vertebrate organisms (Su et al. 2011a; Su et al. 2011b). First, DNA methylation endows genomes with the ability to subject specific sequences to irreversible transcriptional silencing even in the presence of all of the factors required for their expression, an ability that is generally unavailable to organisms that have unmethylated genome (Bestor et al. 2015). Second, DNA methylation may contribute to the functional divergence and retention of duplicated genes (Chang and Liao, 2012; Keller and Yi, 2014). Whether the second round of whole genome duplication in early vertebrates occurred before or after the cyclostome–gnathostome split still remains a controversy (Wang and Gu, 2002; Smith and Keinath 2015). The sea lamprey genome methylation pattern suggests that the characteristic vertebrate global genomic DNA methylation may not have been established in the jawless fishes. Instead, it might have been completely established after the divergence of jawless fishes but before the divergence of teleosts. The high variability of methylation status between individual cells in sea lamprey suggested that the DNA methylation might not be very precisely regulated in early vertebrates, at least before the split of cartilaginous fish. Further investigations of the methylation pattern in cartilaginous fish such as elephant shark will likely contribute to our understanding of the role of WGD in the transition of genome methylation pattern during the evolution of early vertebrates.

DNA methylation patterns are critical to maintain genome stability by repressing transposable elements (TEs) in the genome. The intermediate genome wide methylation level may have contributed to the instability of sea lamprey genome, which contains abundant transposable elements and consists of a lot of small or even tiny microchromosomes (Smith et al. 2013). The sea lamprey may have utilized programmed DNA elimination during early embryogenesis to silence the transposable elements and maintain genome stability (Smith et al. 2012; Haeusser and Margolin 2012). Interestingly, programmed genome change through DNA elimination is also present in some other vertebrates, such as spotted ratfish, bandicoot and zebra finch, and DNA methylation may play an important role in this process (Bracht et al. 2012; Bracht 2014). Similarly, chicken genome also exhibits relatively low genome wide DNA methylation level (Mugal et al. 2015), and contains many tiny microchromosomes (Smith et al. 2013). However, chicken with its characteristic small genome, one of the smallest genome of any terrestrial vertebrate, have repeat contents of only 15~20% (Hillier et al. 2004). This is likely to be a product of an evolutionary process that minimizes the transposable repeats for maintaining genomic integrity and stability to counteract the effects of decreasing of genome methylation level.

Furthermore, we detected differential methylation regions (DMRs) between the three sea lamprey tissues and found a significant DMR between heart and sperm in the promoter of the SEPT9A gene (supplementary figure 8). Because we generated only low-coverage bisulfite sequencing data for sperm and muscle, and the whole genome intermediate methylation level in sea lamprey genome, it is reasonable that only a very small number of DMRs could be detected in our study.

In conclusion, the methylation pattern in sea lamprey provided evidences for the gradual establishment of the vertebrate genomic methylation profile. The dataset in the present study represents the first whole-genome single-nucleotide-resolution methylome of a jawless vertebrate and lays the groundwork for future analyses of the evolution and biological function of DNA methylation in jawless vertebrates.

## Materials and Methods

### Biological materials

All sea lamprey genomic DNA from sperm, heart and muscle was collected from one male lamprey from the Lanrential Great Lakes (Michigan, U.S.A). These tissues were stored in RNAlater before processing. The genomic DNA of the three tissues was purified using the DNeasy blood & tissue kit (QIAGEN). The total RNA was extracted from the same lamprey heart using TRIzol (Invitrogen) according to the manufacturer’s protocol. The quality of the DNA and RNA was monitored using a Nanodrop 2000.

### Whole-genome bisulfite library preparation and sequencing

The libraries were constructed according standard Illumina protocols. We employed bisulfite sequencing to obtain the methylated genomic DNA data for three sea lamprey tissues. Each sample contained 5 μg genomic DNA that was fragmented by Covaris shearing. The fragments were blunt-ended and phosphorylated, and a single A nucleotide was added to the 3′ ends of the fragments in preparation for ligation to a methylated adapter with a single base T overhang. Unmethylated phage lambda genomic DNA (Promega, catalog # D1521) was spiked into each genomic DNA sample as negative control to evaluate the bisulfite conversion efficiency. We also spiked methylated human DNA standard (Zymo Research, catalog # D5011) as positive control to estimate the over-conversion rate for the methylated cytosines. The ligation products were purified and size-selected by agarose gel electrophoresis. The size-elected DNA was treated with bisulfite and purified. The treated DNA was PCR-amplified (8 cycles) to enrich for fragments with adapters on both ends. The final purified products were then quantitated prior to cluster generation. The quality of the libraries and size distribution were assessed using an Agilent 2100 Bioanalyzer (Agilent Technologies). The libraries were sequenced using Illumina HiSeq 2000 machines.

### Genome information

The genome information was downloaded from the ENSEMBL (http://www.ensembl.org) database, including information from the human (version *GRCh37),* western clawed frog *(JGI4.2),* zebrafish *(Zv9),* sea lamprey *(Pmarinus_7.0),* sea squirt *(KH),* and honey bee *(Amel4.0).* The gene annotations were also downloaded from ENSEMBL using consistent versions. The genome and annotation of elephant shark *(Callorhinchus_milii-6.1.3* (Venkatesh et al. 2014)) was obtained from NCBI (http://www.ncbi.nlm.nih.gov.)

### Quality control and sequence alignment

Bisulfite sequencing data for human (GSM752296), zebrafish (GSM1077593), sea squirt (SRR039325) and honey bee (SRP030016) were retrieved from the NCBI database. The downloaded files were decompressed to fastq files using fastq-dump software. Together with the sea lamprey bisulfate sequencing data, all fastq files were examined using FASTQC software (vision 0.10.1, http://www.bioinformatics.babraham.ac.uk/projects/fastqc/) to ensure their quality. Next, high-quality reads were aligned to the appropriate genome using Bowtie2.2.3 (Langmead and Salzberg 2012) available in the Bismark version 0.7.0 (Krueger and Andrews 2011) software packages, allowing up to two mismatches for each paired reads. We filtrated the sites with coverage >5 to perform the following analyses.

### CpG_O/E_ and methylation level estimation

We defined the gene body region using the annotated transcripts, and if multiple transcripts for a gene were present, the longest transcript was used. The region from - 1000 bp to +200 bp of the TSS was defined as the promoter. The 1st exon was also defined based on the annotation information, and for single-exon genes, the +1 bp to +100 bp served as counterparts. The CpG_O/E_ of each gene in the gene body, promoter, exon, intron, and 1st exon regions was defined following previous studies (Elango and Yi 2008; Weber et al. 2007; Suzuki et al. 2007):

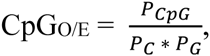

where P_CpG_, P_C_, and P_G_ are the frequencies of the CpG dinucleotides, C nucleotides, and G nucleotides, respectively. The methylation level of each sequenced C was calculated using M/D, M and D strands for the methylated read coverage and total coverage of the C. The average methylation level of the gene body, promoter, exon or intron was defined as follows:

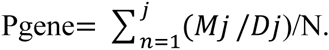

where the M is the methylated read coverage, the D is the total coverage for each C, and N is the total number of C in the specific region. Four regions with a length of 4000 bp upstream or downstream of the TSS or TES, respectively, were used to perform sliding window analyses. We first counted the methylation level of each C using the aforementioned method, and then calculated the methylation level of each specific window (using a previous method). The window size was set as 300 bp, and step was set as 30 bp. The length of the collected regions for the exon-intron-exon and intron-exon-intron structure analyses were 200 bp upstream or downstream of the junction, and the window size and step were 20 and 2 bp, respectively. We used a model based on the binomial distribution to identify the methylation site as previous studies described (Lister et al. 2009), and only the mCs with FDR adjusted P-values < 0.01 were considered true positives.

### Methylation level analyses of transposons

We defined the transposons of each species using the Inverted Repeats Finder (version 3.05) (Warburton et al. 2004) with the following parameters: Match, 2; Mismatch, 3; Delta, 5; PM, 80; PI, 10; Min score, 40. We used the aforementioned sliding window method to analyze the DNA methylation in transposons, with a flanking region length of 3000 bp. The window size and step were set at 200 bp and 20 bp, respectively.

### Analysis of RNA-seq data

The sea lamprey heart mRNA was sequenced as 100 paired-end bases using HiSeq2000. The file for the sea squirt (SRX020114) and zebrafish (SRX209919) RNA-seq reads was downloaded from NCBI and decompressed using fastq-dump. All the fastq files from the three species were evaluated using FastQC (v0.10.1). Bases with a Phred quality score of less than 20 were removed using the fastx toolkit (v0.0.13). All processed reads were mapped to the reference genome using TopHat (v2.0.10) (Trapnell et al. 2009) and bowtie2 (v2.1.0) (Langmead and Salzberg 2012). Multi-mapping reads and duplication reads produced by PCR or poor library construction were filtered out with samtools (v0.1.18) (Li et al. 2009). Then, the unique mapped reads were assembled, and the expression level of a gene was calculated as reads per kilobase of exon model per million mapped reads (FPKM) using cufflinks (v2.1.1) (Trapnell et al. 2010). The expression values were transformed as log2(FPKM+1) for further analysis.

### Homologous genes

We searched the homologous genes in human, zebrafish, sea lamprey and sea squirt in ENSMBL using the BioMart tools with the human genome as a reference. Singlecopy genes of those species were extracted based on one-to-one orthologs.

### Relative DNA Methylation Divergence

We calculated the relative DNA methylation divergence (RDMD) as (M1-M2)/(M1+M2) of paired single-copy genes between every two species, where M1 and M2 are the average methylation levels for the paired genes (Keller and Yi 2014), respectively. The mean RDMD values of all single-copy genes between the two species were calculated as the relative DNA methylation divergence between the two species.

### Phylogenic analyses and domain definition

Human DNMT3A, DNMT3B and DNMT1 protein sequences were obtained from ENSEMBL. Homologous genes in honey bee, sea squirt, sea lamprey and zebrafish were collected using BioMart tools in ENSEMBL. The P-distance was estimated in MEGA6 using default parameters (Tamura et al. 2013). We aligned the protein sequences using MUSCLE (Edgar 2004) and reconstructed the phylogenic tree with PHYML (version 2.1.4) (Guindon et al. 2009) using the GTR+I+G model. Protein domains were predicted by Pfam (http://pfam.xfam.org/). SMART (http://smart.embl-heidelberg.de/) and MEME (http://meme.nbcr.net/meme) were also used to confirm the Pfam predictions.

### DMR definition

We extracted CpG dinucleotides with 5 < q depth < 100 as candidates to detect differentially methylated loci (DML) using the lognormal-beta-binomial hierarchical model. Then, we performed a pairwise comparison of the scaffolds between the three sea lamprey tissues to discover DMRs using the R package DSS in Bioconductor with the Wald test (Feng et al. 2014).

### Data access

The WGBS sequencing data obtained in this study have been deposited in the NCBI Sequence Read Archive (SRA; http://www.ncbi.nlm.nih.gov/sra) under the following accession numbers: SRR2442802, SRR2457525, and SRR2457526. The RNA-seq data from this study have been submitted to the NCBI Gene Expression Omnibus (GEO; http://www.ncbi.nlm.nih.gov/geo/) under accession number GSE73522.

## Acknowledgments

We thank Yu-Wen Chung-Davidson for assistance with sample preparation. This work was supported by grants from the National Natural Science Foundation of China (31272299, 31301034), a grant from the Ministry of Science and Technology China (2012CB910101). Z.S. was supported by Shanghai Pujiang Program (13PJD005).

## Supplementary figure legends

**Figure S1.**
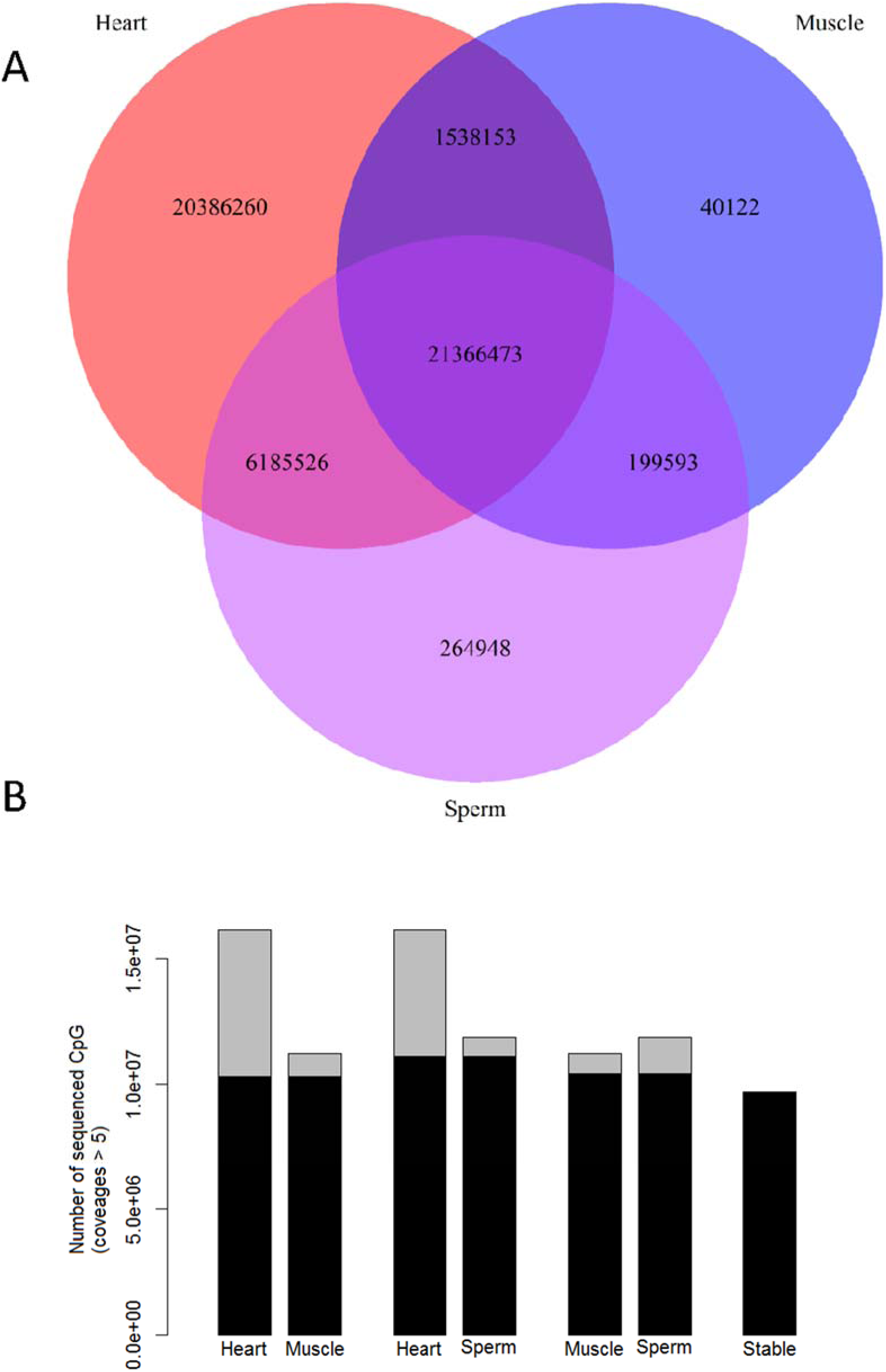
A) Sequencing reads encompassing C in CpGs for three sea lamprey tissues. B) Results for >5x sequenced C in CpGs for three sea lamprey tissues. Black bars share site numbers between/during the three sea lamprey tissues.

**Figure S2.**
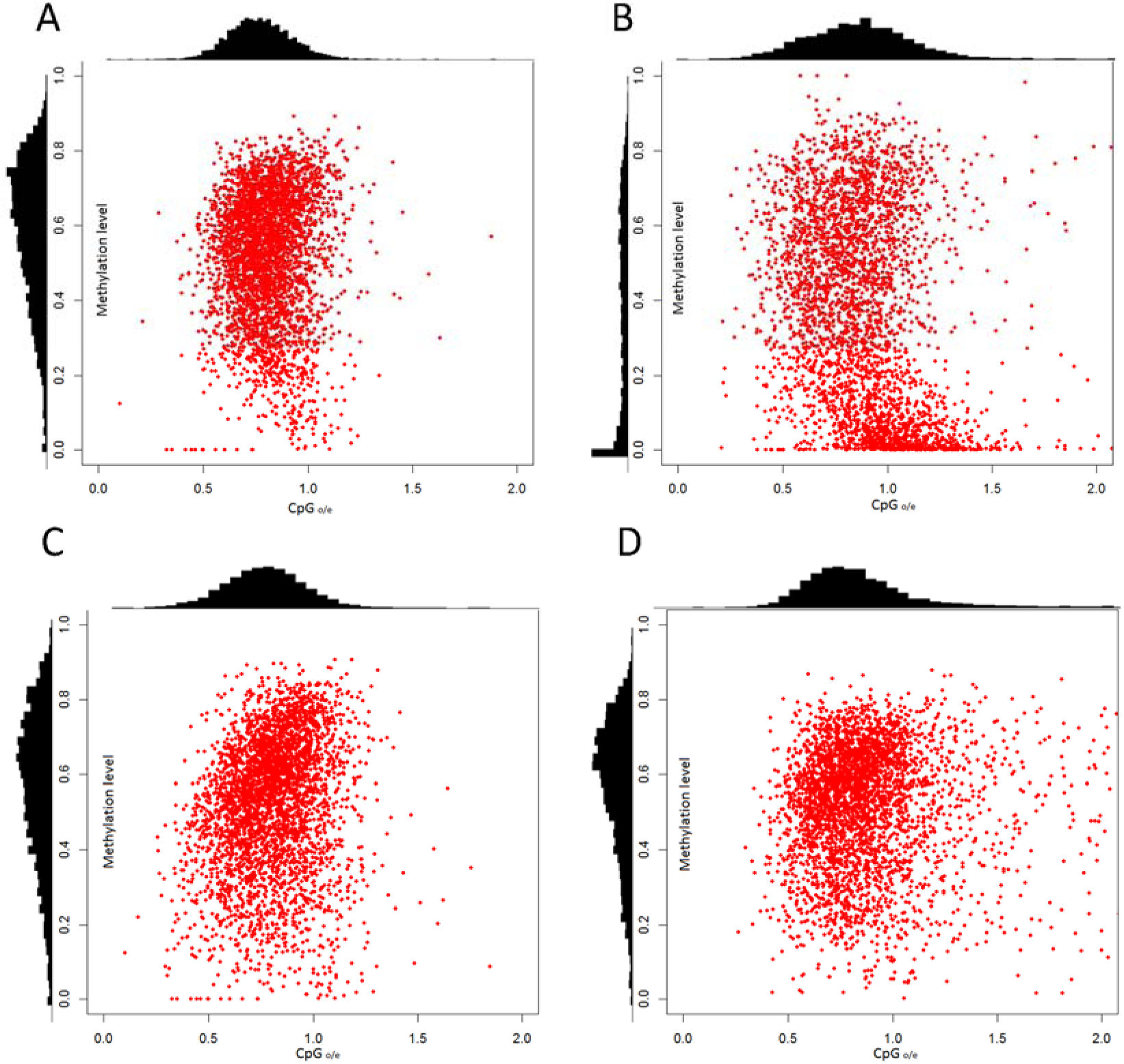
Comparison of CpG O/E ratios and methylation levels in sea lamprey germline genes.

**Figure S3.**
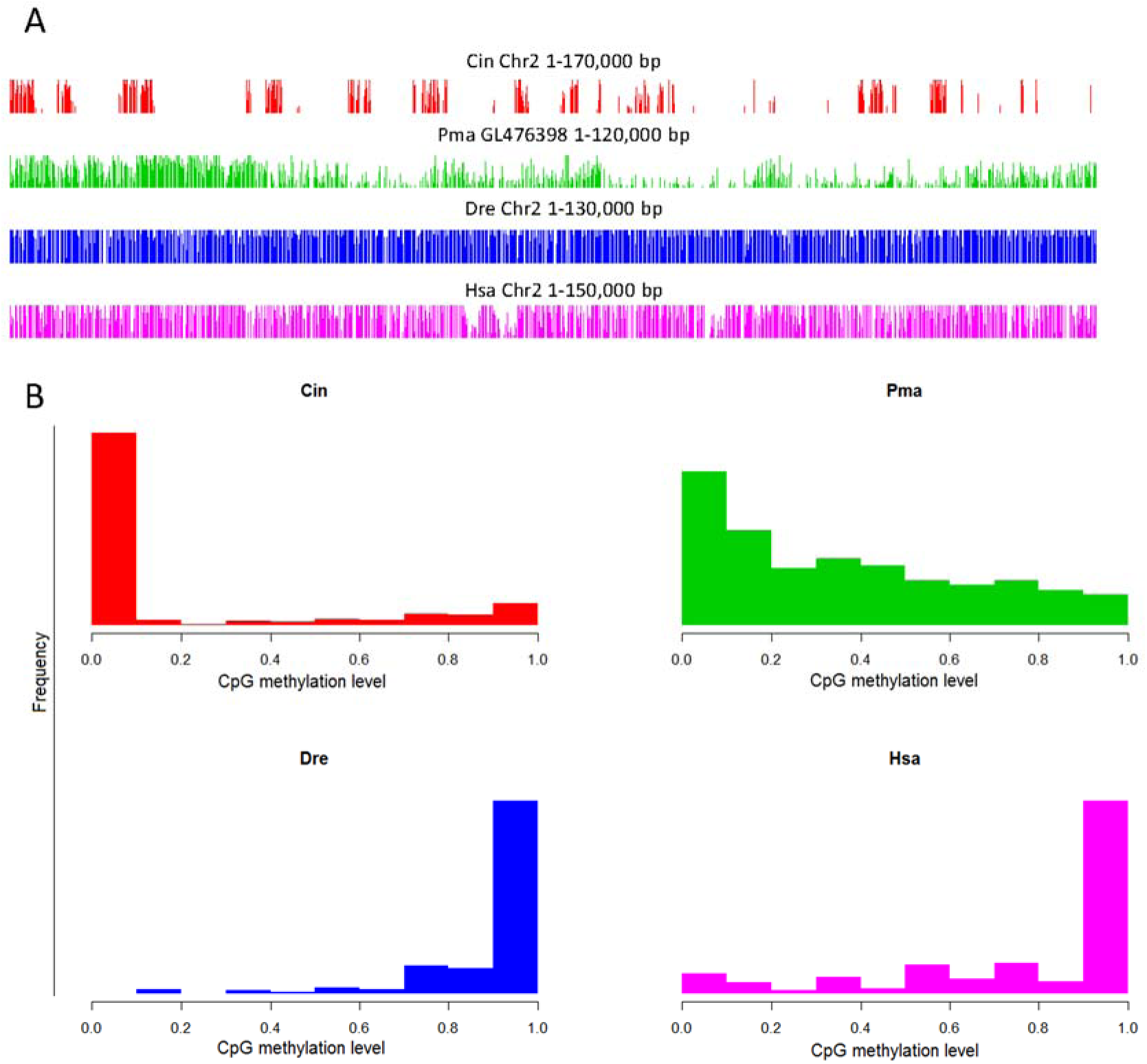
A) DNA methylation patterns determined for a randomly selected genomic region in the four species. B) Distribution of the CpG methylation levels in sea squirt, sea lamprey, zebrafish and human.

**Figure S4.**
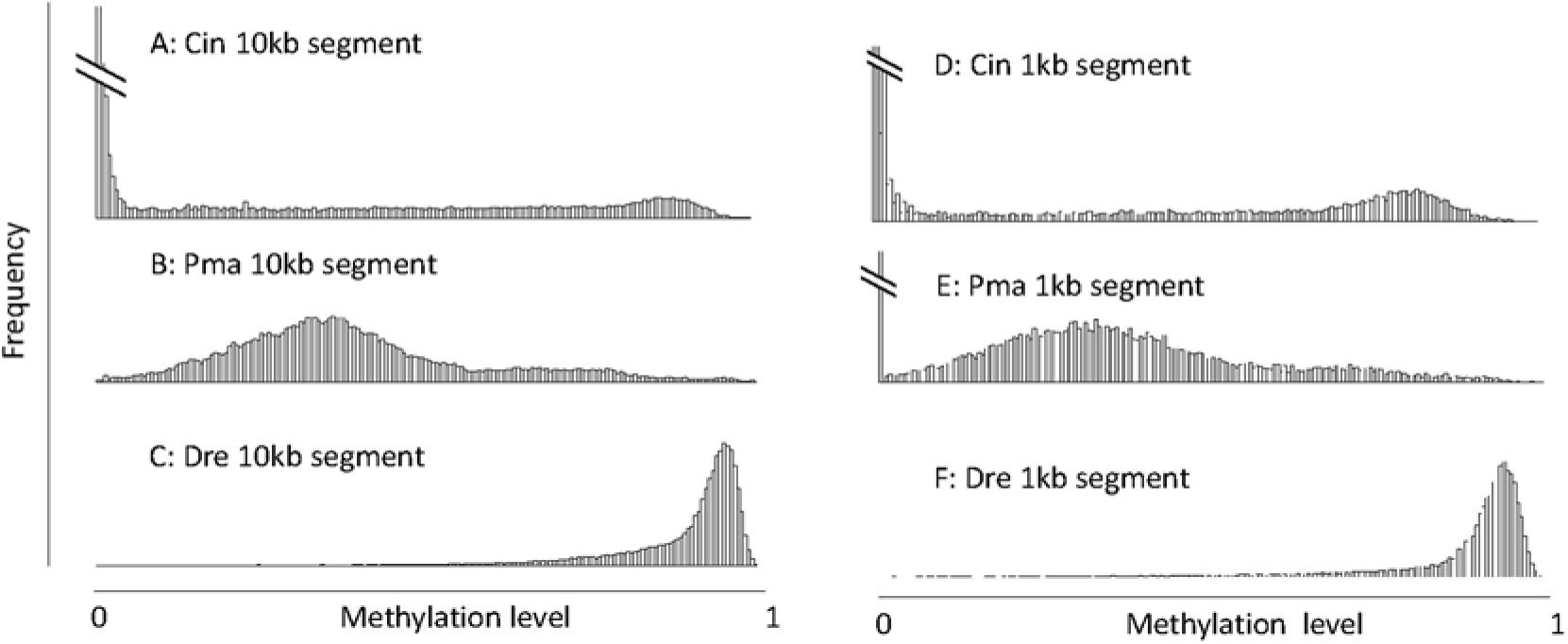
Methylation level distribution of 10kb genomic segments in sea squirt (A), sea lamprey (B), and zebrafish (C) and 1kb genomic segments in sea squirt (D), sea lamprey (E), and zebrafish (F).

**Figure S5.**
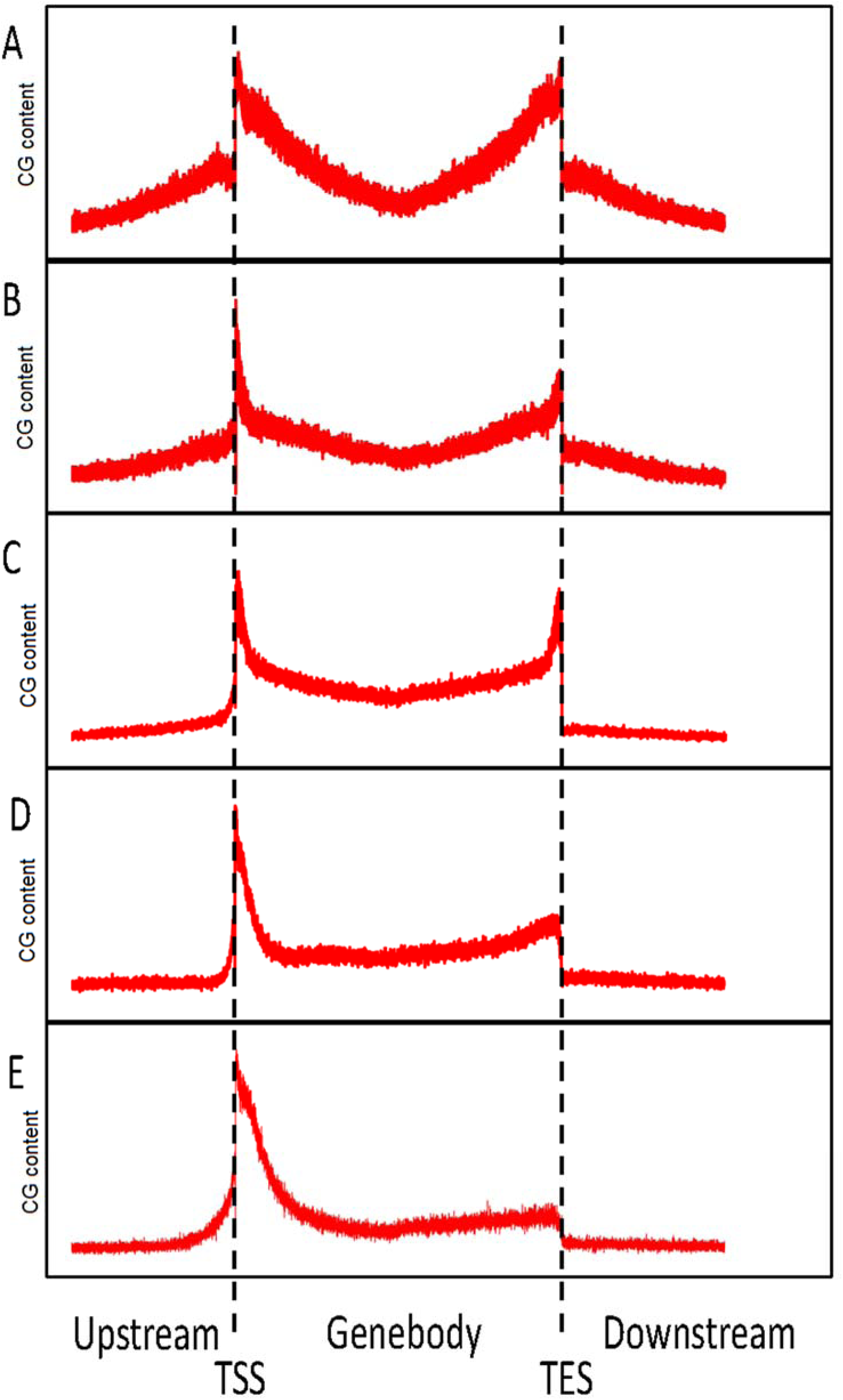
GC content of coding genes and their flanking regions in honey bee (A), sea squirt (B), sea lamprey (C), zebrafish (D) and human (E).

**Figure S6.**
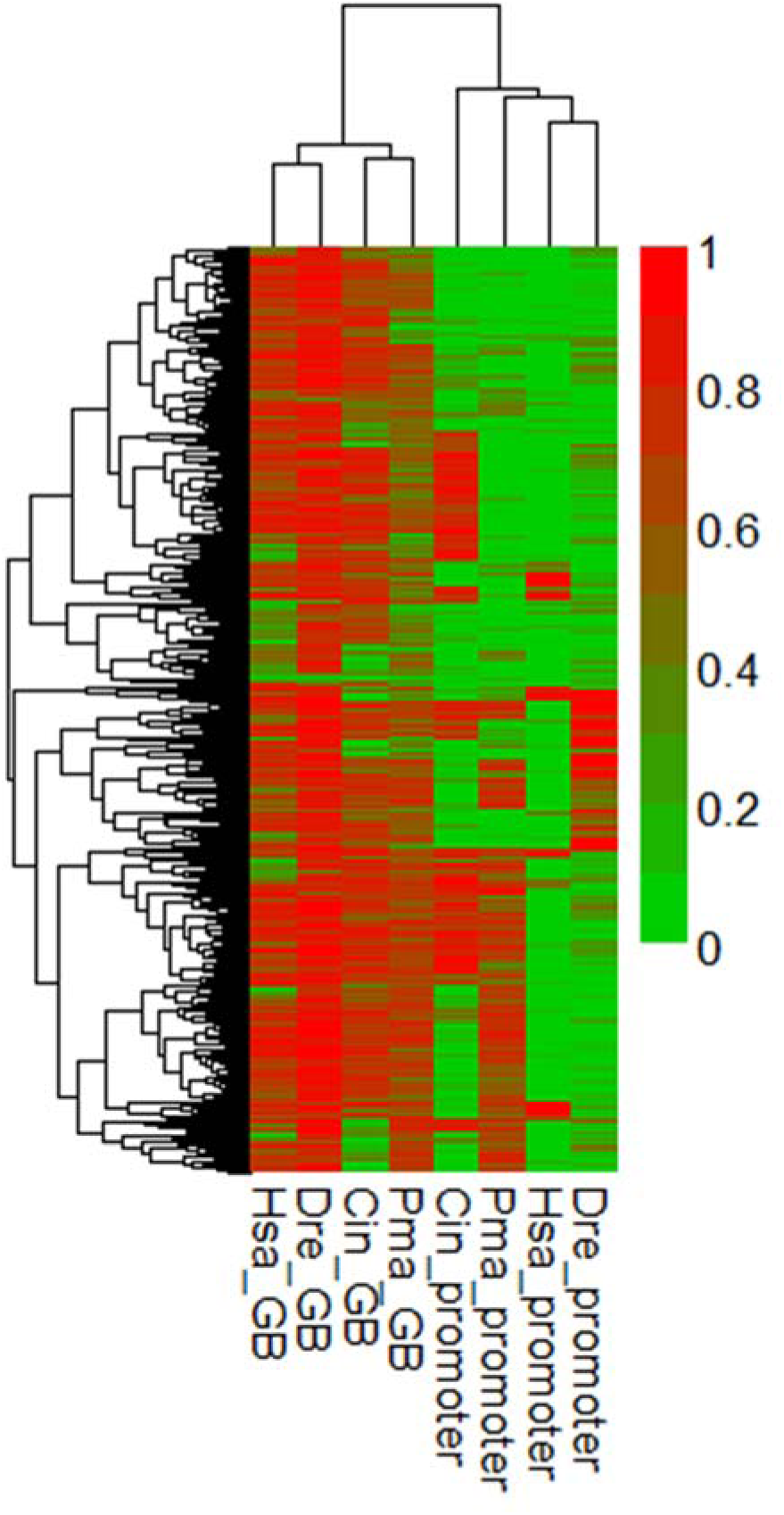
Heat map representations of single-copy gene CpG methylation in gene bodies and promoters in sea squirt, sea lamprey, zebrafish and human. Hierarchical clustering of genes (columns) and categories (rows) was based on 1-r of the methylation level.

**Figure S7.**
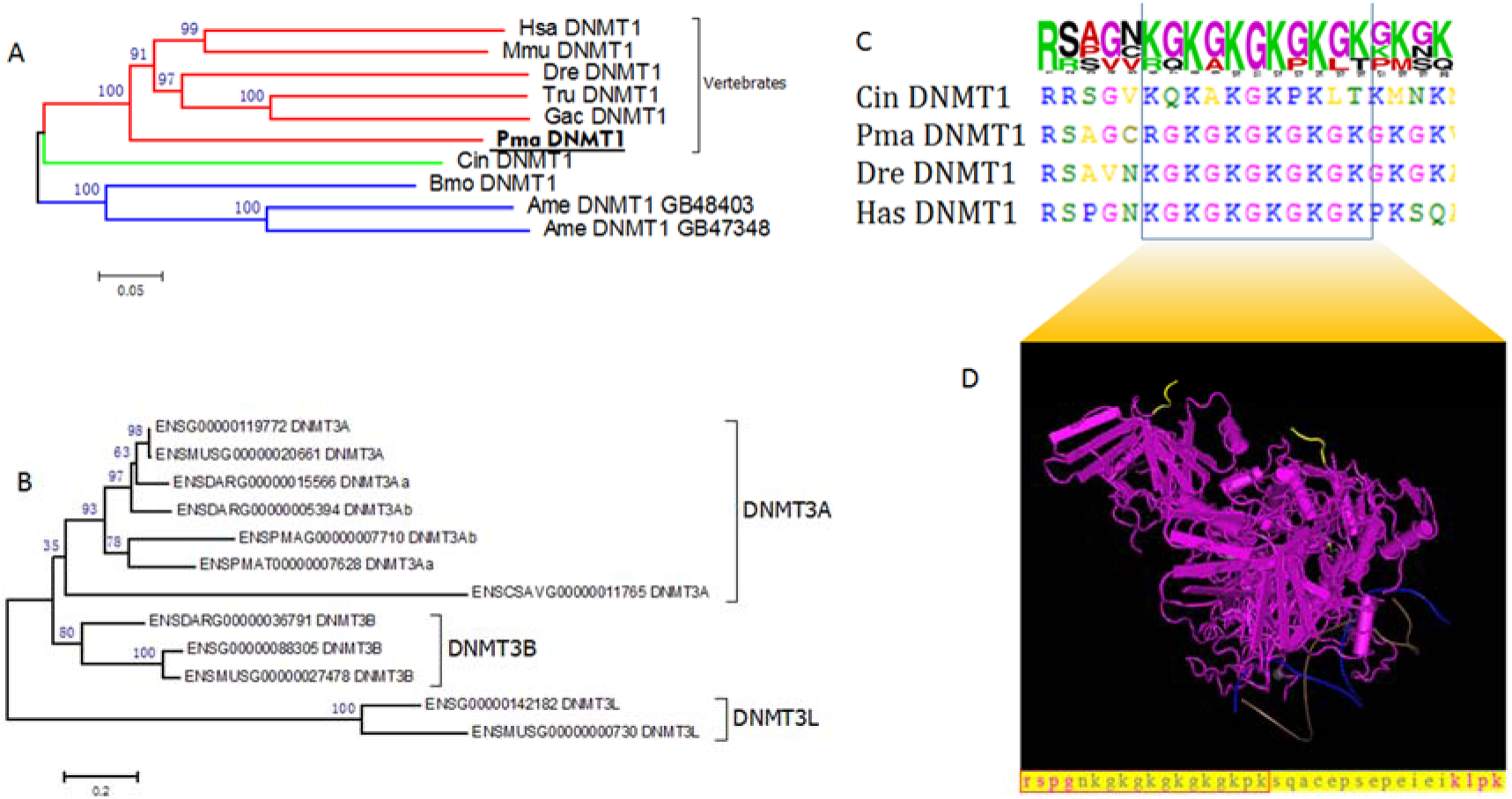
Phylogenetic analyses of DNMT1 (A) and DNMT3 (B). Multiple sequence alignment (C) and 3D structure (D) of the DNMT1 KG-rich motif.

**Figure S8.**
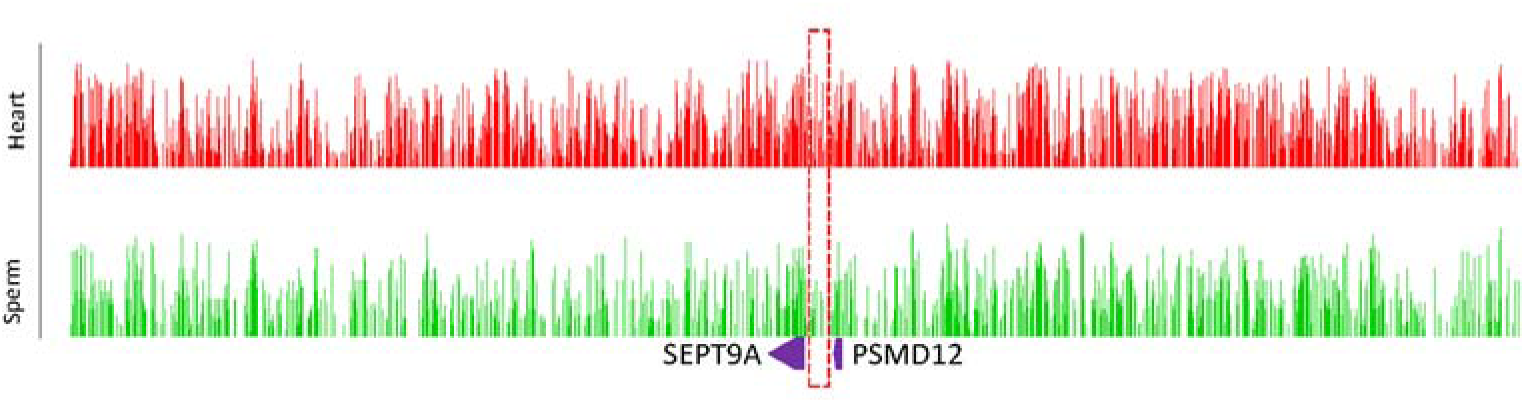
Comparison of the DMRs in the sept9a promoter between sperm and heart.

---

**Supplementary table 1.** Coverage and detail information of the BS-seq in three lamprey tissues

|  |  | *All sites(Mb)* | *Covered sites(Mb)* | *Coverage%* |
| --- | --- | --- | --- | --- |
| *Heart* | CpG | 27.45 | 22.89 | 83.40% |
|  | CHH | 92.79 | 74.04 | 79.80% |
|  | CHG | 28.185 | 23.77 | 84.33% |
| *Sperm* | CpG | 27.45 | 13.59 | 49.50% |
|  | CHH | 92.79 | 31.255 | 33.68% |
|  | CHG | 28.185 | 12.445 | 44.15% |
| *Muscle* | CpG | 27.45 | 7.79 | 28.38% |
|  | CHH | 92.79 | 14.09 | 15.18% |
|  | CHG | 28.185 | 6.75 | 23.95% |

**Supplementary table 2.**
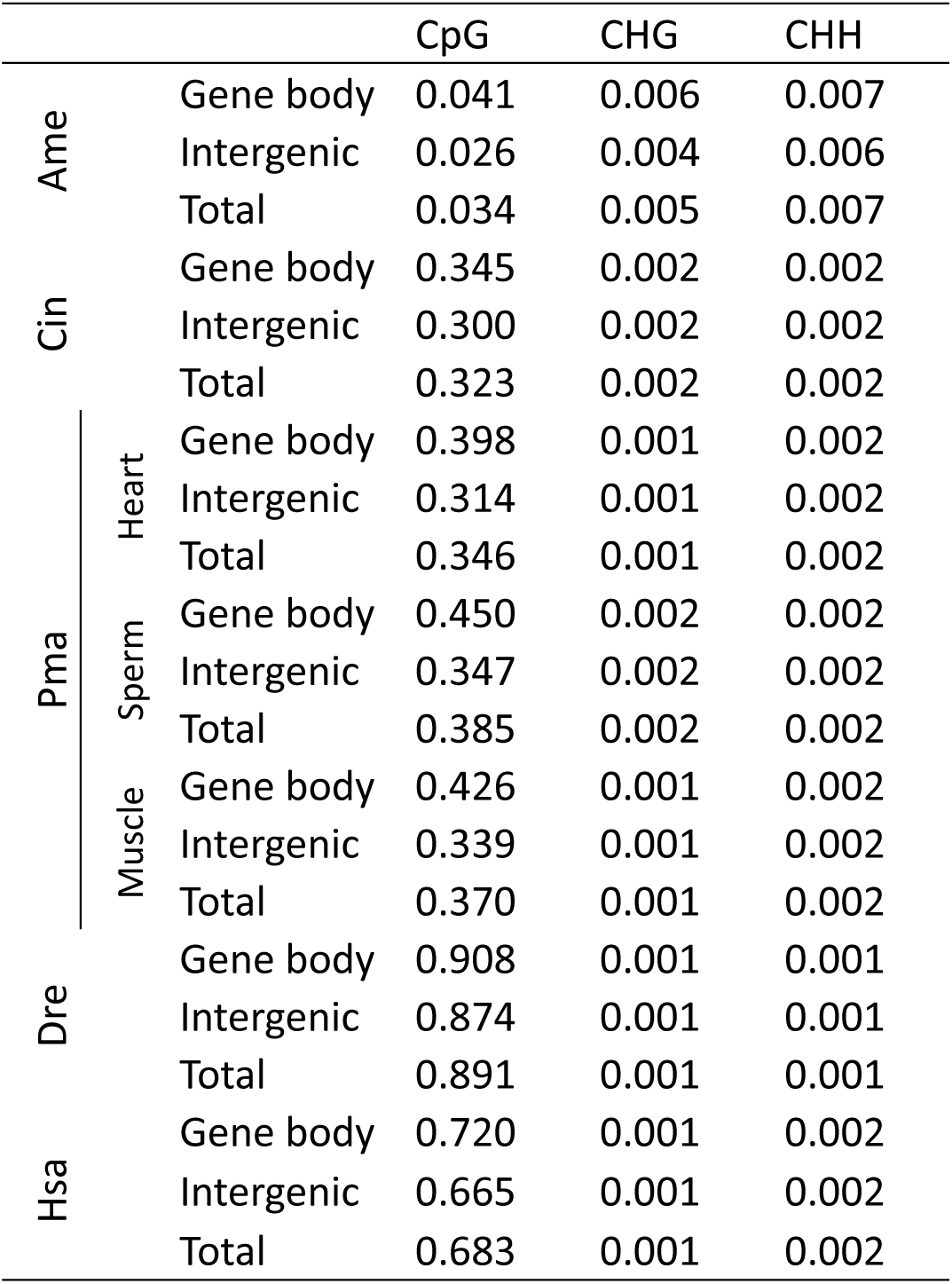
Methylation level of gene body and intergenic region in the five animals and three lamprey tissues.

**Supplementary table 3.** Relative DNA methylation divergence between pairwise species’ single copy genes

|  | Ame | Cin | Pma | Dre | Hsa |
| --- | --- | --- | --- | --- | --- |
| Ame |  | 0.810 | 0.833 | 0.848 | 0.842 |
| Cin | 0.651 |  | **0.072** | 0.121 | 0.101 |
| Pma | 0.743 | **0.180** |  | **0.050** | 0.036 |
| Dre | 0.811 | 0.341 | **0.171** |  | 0.021 |
| Hsa | 0.806 | 0.326 | 0.156 | 0.016 |  |

Up triangle is the promoter and the other is the gene body.

**Supplementary table 4.**
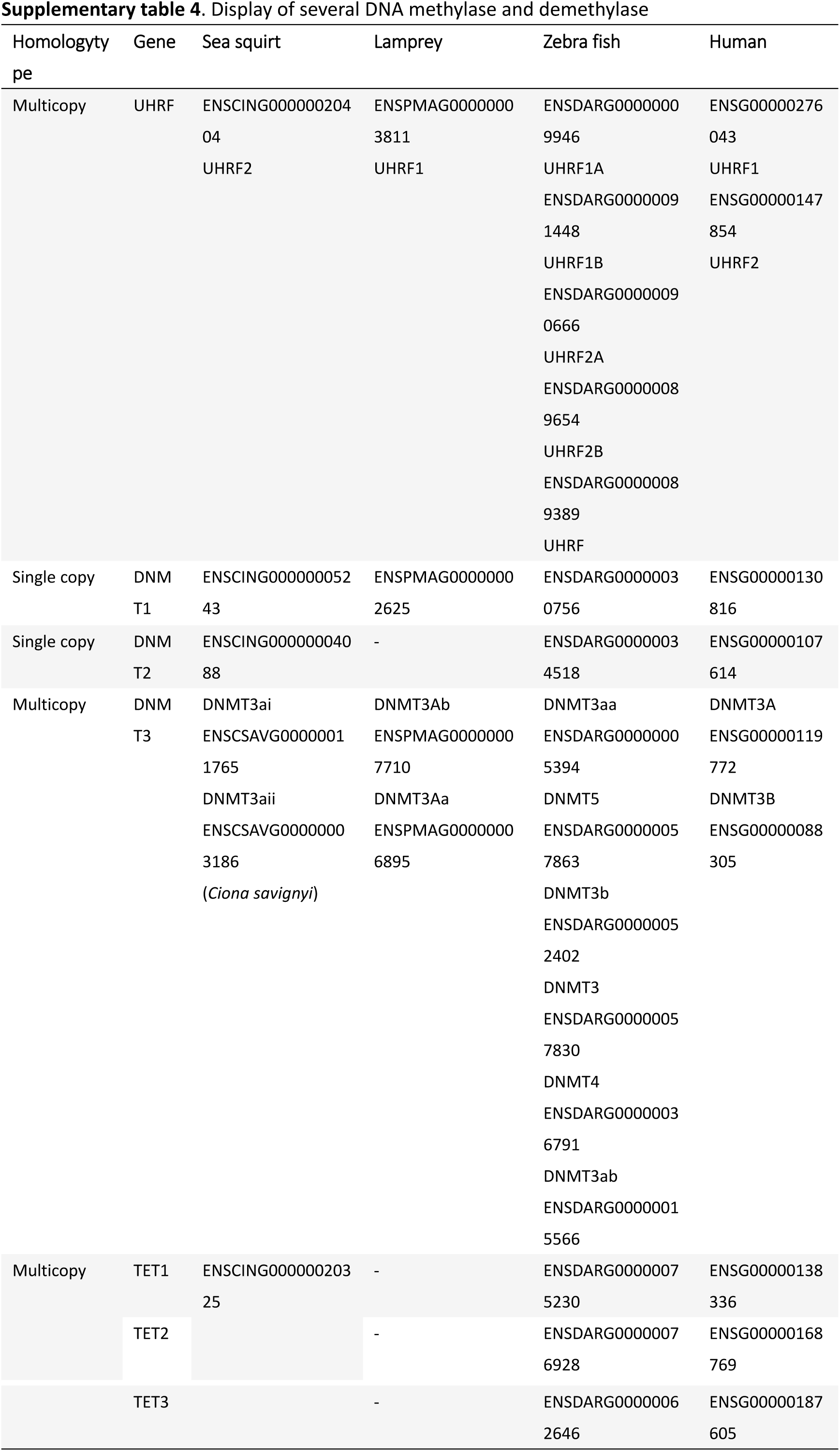
Display of several DNA methylase and demethylase

